# Partitioning variance in cortical morphometry into genetic, environmental, and subject-specific components

**DOI:** 10.1101/2023.07.17.549390

**Authors:** Diana M. Smith, Pravesh Parekh, Joseph Kennedy, Robert Loughnan, Oleksandr Frei, Thomas E. Nichols, Ole A. Andreassen, Terry L. Jernigan, Anders M. Dale

## Abstract

The relative contributions of genetic variation and experience in shaping the morphology of the adolescent brain are not fully understood. Using longitudinal data from 11,665 subjects in the ABCD Study^®^, we fit vertex-wise variance components including family effects, genetic effects, and subject-level effects using a computationally efficient framework. Variance in cortical thickness and surface area is largely attributable to genetic influence, whereas sulcal depth is primarily explained by subject-level effects. Our results identify areas with heterogeneous distributions of heritability estimates that have not been seen in previous work using data from cortical regions. We discuss the biological importance of subject-specific variance and its implications for environmental influences on cortical development and maturation.

## Introduction

The heritability of cortical brain imaging phenotypes has been a subject of investigation for several years. Prior studies have employed both twin datasets (Chen et al. 2013) and genome-wide association studies (GWAS; Yang et al. 2017; Shadrin et al. 2021; van der Meer et al. 2021) to estimate the contribution of genetic variation to variance in cortical morphometry at the region of interest (ROI) and vertex level (Eyler et al. 2012; Chen et al. 2015; Maes et al. 2023). Longitudinal datasets that capture changes in brain structure over time can provide novel insights into the heritability of brain structure. However, until recently, vertex-wise mixed-effects models assessing the influence of shared genetic variance, as well as family- and subject-specific variance, on cortical morphometry have not yet been available, due in part to the computational demand of running complex models on tens of thousands of cortical vertices. Here we used a novel computational method to apply mixed-effects models to the large sample and the longitudinal design of the Adolescent Brain Cognitive Development⍰ Study (ABCD Study^®^), to estimate the contribution of genetic relatedness (heritability), shared family environment, and subject-specific variance to vertex-wise measures of brain morphometry.

To examine the spatial distributions of random effects across vertex-wise cortical measures, we applied a novel method, Fast and Efficient Mixed-Effects Algorithm (FEMA; Parekh et al. 2023), which can model genetic relatedness as a continuous value ranging from 0 to 1, rather than assigning categorical variables based on kinship. FEMA also allows for the flexible specification of several random effects simultaneously including shared family, subject ID, and others(Parekh et al. 2023). By using the full ABCD Study^®^ sample (n=11,880) rather than restricting our analyses to twins, we were able to better approximate the variance components that exist in a general population. This approach to heritability analysis also has the potential to capture more shared variance than the heritability estimates obtained from GWAS, which only model the additive effects that can be inferred from common SNPs and therefore may not capture variance attributable to structural variants, rare variants, or non-additive effects (Génin 2020). In addition, incorporation of genetic, family, and subject-specific effects in the same model allows us to investigate the differential effects of genetics, shared environment, and otherwise unexplained contributions to within-subject stability.

We estimated the contribution of additive genetic relatedness (akin to a heritability estimate), shared family environment (the extent to which siblings and twins share variance, coded as shared family ID), and the effect of subject (variance that is *not* explained by fixed effect covariates nor genetic/family structure, but nevertheless remains stable for a given subject over time). Of note, classical additive genetic / common environment / unique environment (ACE) models typically include data from a single timepoint and therefore cannot inspect the contribution of subject-specific variance in this manner. Using longitudinal neuroimaging data from the ABCD Study^®^ release 4.0, we derived vertex-wise cortical thickness, cortical surface area, and sulcal depth. Then, for each vertex, we used the FEMA package (Parekh et al. 2023) to fit the model

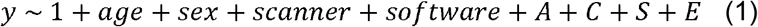

where *y* represents the phenotype at each vertex; age, sex, MRI scanner, and scanner software version are fixed effect covariates; *A, C, S*, and *E* are the random effects, with their estimates corresponding to the proportion of variance in the phenotype (not explained by fixed effects) attributable to genetic similarity (*A*, modeled as the genetic relatedness between each pair of individuals within the same family, computed from SNP data), common family environment (*C*, coded as family ID), subject (*S*), and the remaining unexplained variance (*E*), respectively. Of note, the contribution of the random effect of subject (*S*) is equivalent to the intra-class correlation coefficient (ICC) estimated using a mixed effects model to measure test-retest reliability (Zuo and Xing 2014). Thus, a high value for *S* denotes a phenotype with high test-retest reliability, high values for *A* reflect phenotypes that are highly heritable, and high values for *C* reflect phenotypes that are highly attributable to shared family environment.

## Materials and Methods

### Sample

The ABCD Study^®^ is an ongoing longitudinal multisite study within the United States that includes data from 11,880 adolescents recruited from 21 data acquisition sites (Garavan et al. 2018; Volkow et al. 2018). Each site obtained approval from their Institutional Review Board, and all participants and caregivers underwent verbal and written consent/assent. Exclusion criteria for the ABCD Study^®^ include 1) lack of English proficiency in the child; 2) the presence of severe sensory, neurological, medical or intellectual limitations that would inhibit the child’s ability to comply with the study protocol; 3) an inability to complete an MRI scan at baseline. In this study, we used baseline and the first follow-up imaging data (acquired two years after the baseline) from ABCD release 4.0. We included all individuals with imaging and genomics data that passed quality control; analyses were restricted to observations with complete imaging and covariate data (final N = 11,402 at baseline, 7,695 at 2 year follow-up, for a total of 19,097 observations; mean age at first visit = 9.92 years (SD = 0.62), mean age at second visit = 11.94 years (SD = 0.65)). Table 1 shows the demographics of the analytic sample at baseline and at the follow up.

**Table 1.** Demographic data for age in months (mean, (SD)), sex at birth, household income, parental education, parental marital status, self-declared race, and endorsement of Hispanic ethnicity, stratified by time point (baseline and 2-year follow-up). Variable names from the tabulated data release are included in the table for replication.

|  | Baseline | 2 Year Follow-Up | p |
| --- | --- | --- | --- |
| n | 11402 | 7695 |  |
| Number of Families | 9529 | 6524 |  |
| interview_age (years), mean (SD) | 9.92 (0.63) | 11.94 (0.65) | <0.001 |
| sex = M (%) | 5931 (52.0) | 4129 (53.7) | 0.027 |
| household.income (%) |  |  | <0.001 |
| [<50K] | 3074 (29.5) | 1718 (24.2) |  |
| [>=50K & <100K] | 2949 (28.3) | 2006 (28.3) |  |
| [>=100K] | 4409 (42.3) | 3374 (47.5) |  |
| high.educ (%) |  |  | 0.026 |
| < HS Diploma | 560 (4.9) | 334 (4.4) |  |
| HS Diploma/GED | 1078 (9.5) | 677 (8.8) |  |
| Some College | 2950 (25.9) | 1908 (24.9) |  |
| Bachelor | 2902 (25.5) | 2022 (26.3) |  |
| Post Graduate Degree | 3898 (34.2) | 2734 (35.6) |  |
| married = Yes (%) | 7710 (68.2) | 5251 (68.8) | 0.405 |
| race.4level (%) |  |  | 0.004 |
| White | 7267 (64.7) | 5075 (66.8) |  |
| Black | 1754 (15.6) | 1059 (13.9) |  |
| Asian | 263 (2.3) | 157 (2.1) |  |
| Other/Mixed | 1950 (17.4) | 1307 (17.2) |  |
| hisp = Yes (%) | 2332 (20.7) | 1487 (19.6) | 0.054 |

### Genotyping, Genetic Principal Components and Genetic Relatedness

Methods for collecting genetic data have been described in detail elsewhere (Uban et al. 2018). Briefly, a saliva sample was collected at the baseline visit, as well as a blood sample from twin pairs. The Smokescreen™ Genotyping array (Baurley et al. 2016) was used to assay over 500,000 single nucleotide polymorphisms (SNPs), which were used for genetic relatedness calculation using PC-Air (Conomos et al. 2015) and PC-Relate (Conomos et al. 2016). PC-AiR captures ancestry information that is not confounded by relatedness by finding a set of unrelated individuals in the sample that have the highest divergent ancestry and computes the PCs in this set; the remaining related individuals are then projected into this space. PC-Relate computes a GRM that is Independent from ancestry effects as derived from PC-AiR. PC-AiR was run using the default suggested parameters from the GENESIS package (Gogarten et al. 2019), as described in previous work (Smith et al. 2023). Supplementary Figure 6 displays a histogram of the pairwise genetic relatedness values across the full sample (Supp. Fig. 6A) as well as the subset of pairs of participants with shared family ID (Supp Fig. 6B). Of the 2011 pairs of individuals that shared a family ID, 1868 pairs had genomic relatedness data; of these, 1378 pairs had genetic relatedness between 0.25 and 0.75 (most likely full siblings or dizygotic twins) and 389 pairs had genetic relatedness greater than 0.75 (most likely monozygotic twins).

### MRI acquisition and image processing

The ABCD Study^®^ MRI data were collected across 21 research sites using Siemens Prisma, GE 750 and Philips Achieva and Ingenia 3 T scanners. Scanning protocols were harmonized across sites. Details of imaging acquisition and processing protocols used in the ABCD Study^®^ have been described previously (Casey et al. 2018; Hagler et al. 2019). Briefly, T1-weighted images were acquired using a 3D magnetization-prepared rapid acquisition gradient echo (MPRAGE) scan with 1 mm isotropic resolution and no multiband acceleration. T1w structural images were corrected for gradient nonlinearity distortions using scanner-specific, nonlinear transformations provided by MRI scanner manufacturers (Wald et al. 2001; Jovicich et al. 2006). Intensity inhomogeneity correction was performed by applying smoothly varying, estimated B1-bias field (Hagler et al. 2019). Images were rigidly registered using a cross-sectional framework and resampled into alignment with a pre-existing, in-house, averaged, reference brain with 1.0 mm isotropic resolution (Hagler et al. 2019). Cortical surface reconstruction was conducted using FreeSurfer v7.1.1, which includes tools for estimation of various measures of brain morphometry and uses routinely acquired T_1_w MRI volumes (Dale and Sereno 1993; Dale et al. 1999; Fischl, Sereno, and Dale 1999; Fischl, Sereno, Tootell, et al. 1999; Fischl and Dale 2000; Fischl et al. 2001, 2002, 2004; Ségonne et al. 2004, 2007). The cortical parcellation used in this analysis was conducted in FreeSurfer using the Desikan-Kiliany cortical atlas (Desikan et al. 2006). All analyses included only those participants who were recommended for inclusion in post-processed sMRI quality control (imgincl_t1w_include==1).

### Statistical analysis

The classic ACE model used to estimate heritability is equivalent to a linear mixed-effects (LME) model specified as follows:

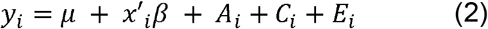

where *y*_*i*_ is the trait value of the *i*^th^ scan; μ is the overall mean; *x*_*i*_ denotes a vector of covariates; and *A*_*i*_, *C*_*i*_, *E*_*i*_ represent latent additive genetic, common family and unique environmental random effects, respectively. Over subjects, the covariance of these three terms are 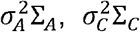, and 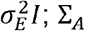 is given by the genetic relatedness, which could be a kinship coefficient (e.g. ½ for siblings or dizygotic twins) or, as we have done, the SNP-wise genetic similarity similar to previous methods (Yang et al. 2011); *Σ*_C_ has 1’s on off-diagonals for any family pairs, 0 otherwise.

For longitudinal datasets incorporating data from multiple timepoints for a given participant, an additional random effect *S* can be incorporated:

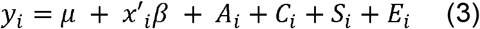

where *S*_*i*_ is the random effect of subject (e.g., subject ID), and the vector of covariates *x*_*i*_ includes a fixed effect to incorporate multiple timepoints (e.g., age). The covariance of the subject effect is 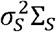, where Σ_*s*_ has 1’s on off diagonals observation pairs from the same subject; with a subject effect modeled, the final effect *E*_i_ corresponds to a pure intrasubject measurement error. Note that the total residual variance, 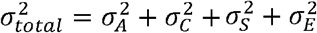, is the same variance that would be estimated in a cross-sectional analysis of unrelated subjects, and demonstrates how this approach decomposes all phenotypic variance into meaningful components.

### Vertex-wise analysis

Univariate linear mixed effects models (LMMs) were applied at each vertex to model cortical morphometry (cortical thickness, cortical surface area, sulcal depth) as the dependent variables.

All of the results shown are from a model including age, sex, MRI scanner and software version as fixed covariates. Random effects were modeled as genetic relatedness (A) and subject (S) nested within shared family groups (C). All LMMs were run using the publicly available FEMA software package (Parekh et al. 2023), which handles voxel- and vertex-wise data and can incorporate a matrix of SNP-derived genetic relatedness.

For the main results, vertex-wise statistical maps present *σ*^2^, the proportion of residual variance that is explained by variance in the random effect of interest. Unthresholded *σ*^2^ maps are presented in the main figures to provide a comprehensive description of the continuous distribution of effects. Supplementary Figure 1 displays the total residual variance, 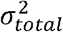, which is the total unexplained variance after accounting for the fixed effects in the model (age, sex, scanner, and software). 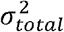 is displayed in units that match the units of the phenotype of interest (mm or mm^2^) and represents the total phenotypic variance that is then partitioned into *A, C, S*, and *E* components.

### Region-of-interest (ROI) analyses

To visualize the distribution of test statistics by region of interest (ROI), the vertex-level test statistics were mapped to the corresponding regions of interest (ROIs) in the Desikan-Kiliany 40 Atlas (Desikan et al. 2006). Violin plots were generated to show the distribution of vertex-level effects across all vertices within each ROI mask, stratified by hemisphere, to highlight the range of effects within each ROI.

All statistical analyses were conducted using custom code in MATLAB v2020a. FEMA is publicly available on GitHub (https://github.com/cmig-research-group/cmig_tools).

## Results

Supplementary Figure 1 presents the total residual variance, *σ*^2^ _*total*_, for cortical thickness, cortical surface area, and sulcal depth. The total residual variance *σ*^2^_*total*_ represents the phenotypic variance that is unexplained by the fixed effects included in our model; *σ*^2^_*total*_ is partitioned into *A, C, S*, and *E* components. Cortical thickness had a relatively uniform distribution of total residual variance, with the largest variances occurring in the temporal pole. Cortical surface area exhibited the largest residual variances in the lateral frontal and occipital cortices, with notably lower residual variances occurring in the primary motor cortex, primary sensory cortex, medial frontal, and medial temporal cortices. Sulcal depth exhibited the largest range in total residual variance, with most regions exhibiting relatively smaller variances whereas the superior parietal lobule exhibited larger residual variances bilaterally. Regions with lower residual variances may represent portions of the cortex that are genetically conserved, whereas regions with higher residual variance may be prone to exhibiting individual differences based on genetic and environmental factors.

Figure 1 presents vertex-wise variance component estimates for cortical surface area as a fraction of total residual variance, *σ*^2^_*total*_. The shared environment (*C*) component accounted for a small proportion of residual variance across the whole brain (Fig. 1), mostly limited to the bilateral temporal poles, medial frontal and occipital cortices, and parts of the primary motor and primary somatosensory cortices. Supp. Fig. 2 displays the distribution of random effects estimates within each region of interest (ROI); *C* estimates ranged from 0.11 (supramarginal gyrus) to 0.26 (temporal pole; Supp. Fig. 2). The *A* component accounted for a larger proportion of variance, with the strongest contributions in the medial frontal and occipital cortices, as well as the superior frontal gyrus and subregions of the superior temporal and insular cortices (Fig. 1). When grouping vertices by ROI, mean *A* component estimates ranged from 0.28 (entorhinal cortex) to 0.57 (pericalcarine cortex; Supp. Fig. 2). Subject-specific variance *S* accounted for the largest proportion of variance in cortical surface area across several regions that were not clearly circumscribed by atlas parcellations, including parts of the posterior cingulate, supramarginal, superior parietal and inferior parietal cortices, reflecting that there is a substantial amount of variance in cortical surface area that is unexplained by genetic or common environmental factors that nonetheless remains stable within subjects over time. After grouping vertices into ROIs, the mean *S* estimates ranged from 0.19 (rostral anterior cingulate cortex) to 0.51 (inferior parietal cortex; Supp. Fig. 2). Inspection of the distribution of random effects estimates within each region showed evidence for right-left asymmetry in some regions (e.g., the banks of the superior temporal sulcus, the entorhinal cortex, *A* and *C* components in the parahippocampal cortex, and *C* in pars orbitalis and pars triangularis). In addition, some parcellated regions exhibited wide distributions of vertex-level effect estimates (e.g., posterior cingulate, precentral, superior frontal, superior parietal, and superior temporal cortices), reflecting the heterogeneity within these regions, whereas others exhibited narrower distributions (e.g., medial orbitofrontal cortex, pars opercularis, pericalcarine cortex, and rostral anterior cingulate cortex).

**Figure 1.**
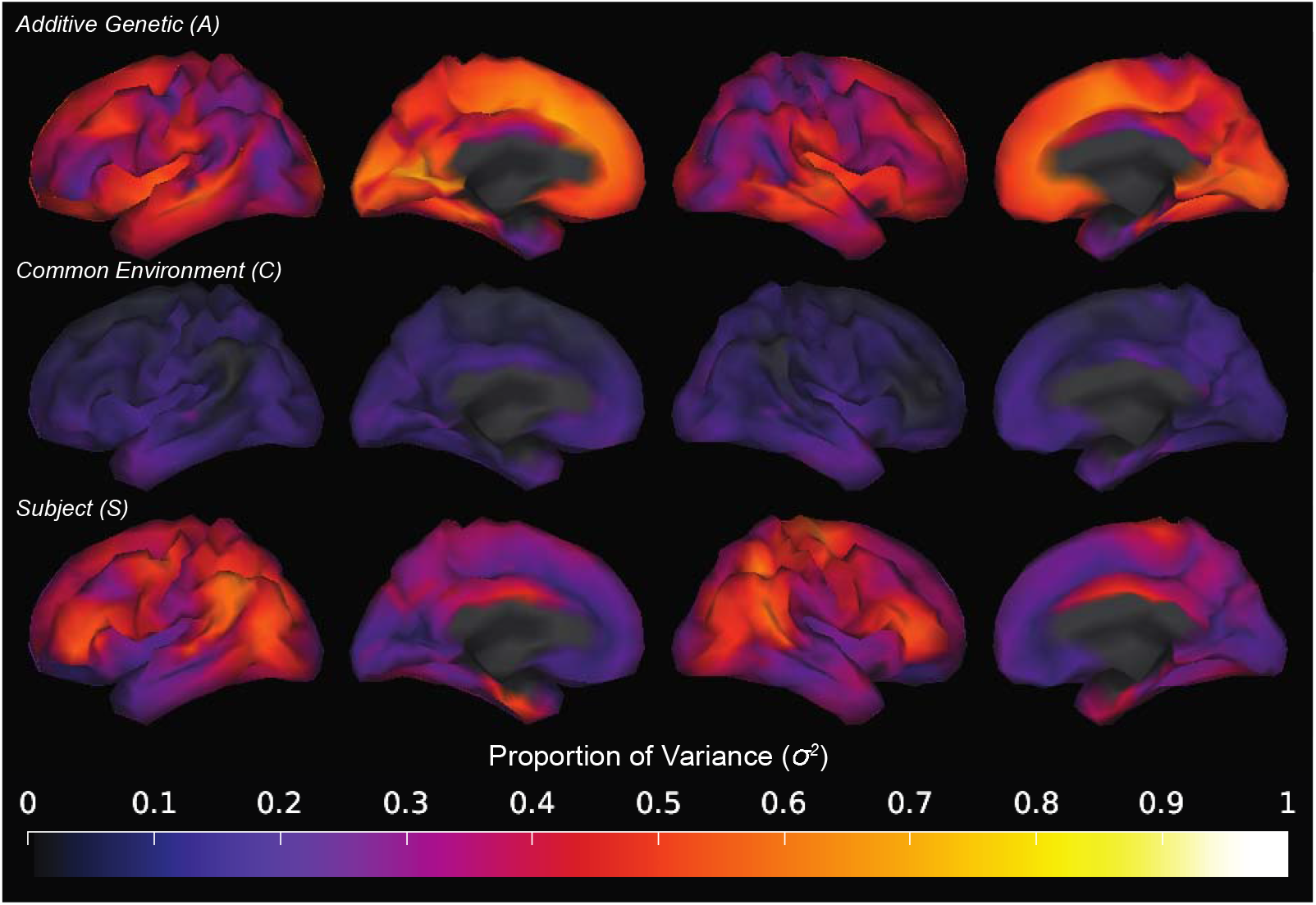
Additive genetic (A), common environment (C), and subject-specific variance (S) components in cortical surface area, presented vertex-wise as a fraction of total residual variance.

Figure 2 presents vertex-wise variance component results for cortical thickness; ROI-wise results are presented in Supp. Fig. 3. The *A* component accounted for a large proportion of residual variance, with particularly strong effects in subregions of the superior frontal and pericalcarine cortices as well as the cuneus, and very small areas in the precentral cortex and insula (Fig. 2). After grouping vertices into ROIs, the mean *A* component estimates across all vertices within an ROI ranged from 0.19 (temporal pole) to 0.44 (cuneus; Supp. Fig. 3). The *S* component also accounted for a substantial proportion of variance in several areas, particularly in the isthmus of the left cingulate cortex and a larger portion of the right cingulate cortex, as well as portions of the superior parietal cortex and the precuneus (Fig. 2). After grouping vertex-level estimates into ROIs, mean *S* estimates across vertices within ROIs ranged from 0.25 (pericalcarine cortex) to 0.42 (isthmus of cingulate cortex; Supp. Fig. 3). Similar to cortical surface area, the *C* component accounted for relatively smaller proportions of residual variance across the cortex, with the highest estimates occurring in areas that also exhibited strong *S* effects (e.g, precuneus and parts of superior parietal cortex; Fig. 2). At the ROI level, mean estimates for the *C* component ranged from 0.09 (pars opercularis) to 0.17 (lingual cortex; Supp. Fig. 3). As with cortical surface area, there were regions that exhibited right-left asymmetry (e.g. *A* estimates in the pars opercularis and pericalcarine cortex) as well as wide or irregular distributions of vertex-level estimates (e.g., *S* estimates in the supramarginal and transverse temporal cortices).

**Figure 2.**
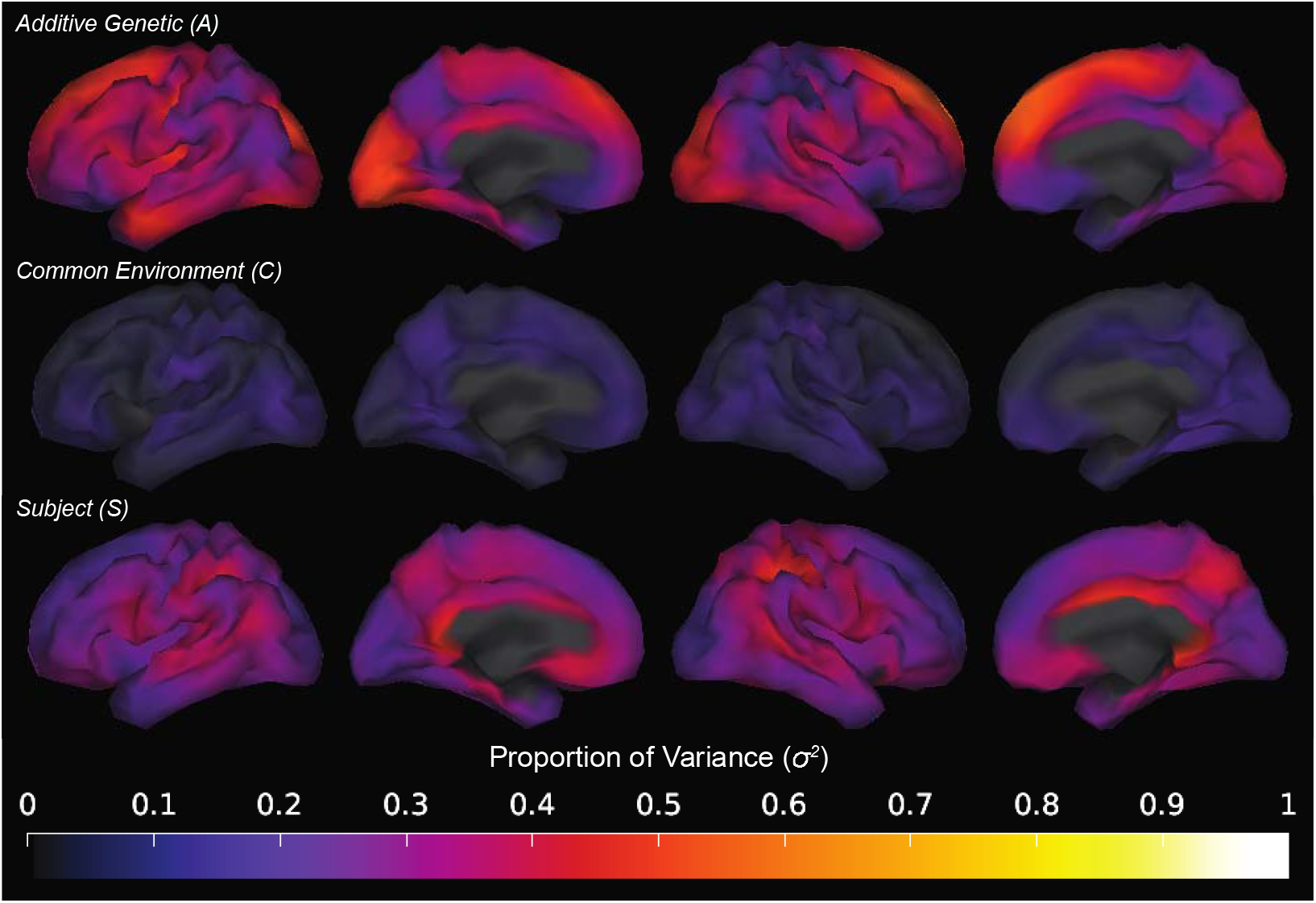
Additive genetic (A), common environment (C), and subject-specific (S) variance components in cortical thickness, presented vertex-wise as a fraction of total residual variance.

Figure 3 presents vertex-wise variance component estimates for sulcal depth. Similar to cortical surface area and thickness, the *C* component accounted for a very small proportion of variance across the whole brain. Unlike the prior phenotypes, however, the *A* effect estimate was also relatively small across most of the brain, with substantial effects limited to portions of the posterior cingulate cortex and isthmus of the cingulate cortex, as well as a very anterior subregion of the precuneus. Conversely, *S* accounted for a large proportion of variance across the entire cortex, with variance component estimates ranging from 0.34 (temporal pole) to 0.74 (caudal middle frontal gyrus; Supp. Fig. 4). Areas with lower *S* estimates were mainly limited to the insula and parts of the entorhinal cortex. Once again, we observed regions with apparent right-left asymmetry (e.g., lingual cortex, pars triangularis, posterior cingulate cortex), though this phenomenon was most apparent for the estimates of the *A* random effect. We also observed wide and/or irregular distributions of vertex level effects within several regions (e.g., A estimates in the precuneus and rostral middle frontal cortex, and S estimates in the supramarginal cortex). Supplementary Figure 5 presents the *A, C*, and *S* variance components in a single summary figure using a red-green-blue color map for ease of interpretation.

**Figure 3.**
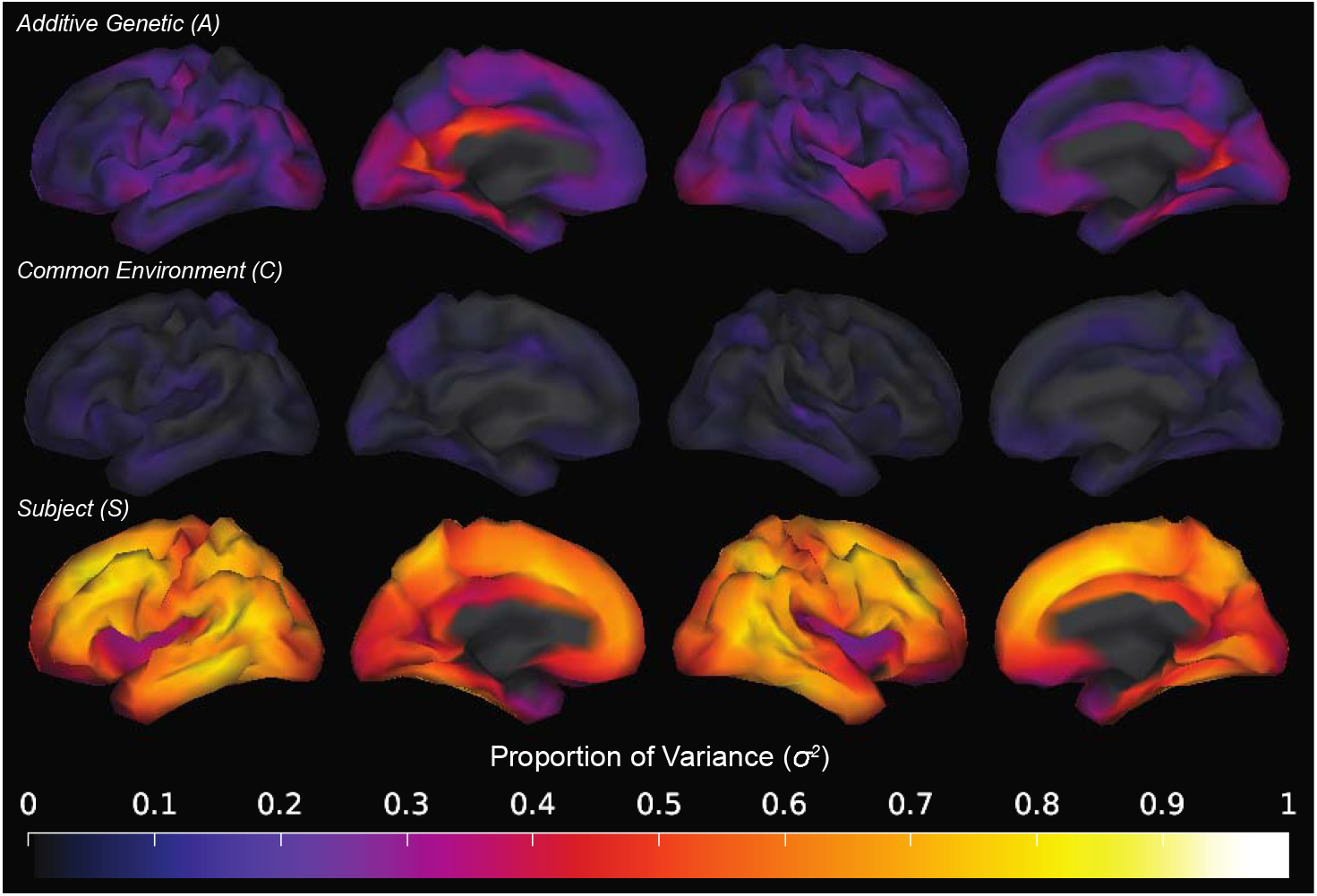
Additive genetic (A), common environment (B), and subject-specific (C) variance components in sulcal depth, presented vertex-wise as a fraction of total residual variance.

## Discussion

Our analysis extends previous investigations of heritability based on twin data (Maes et al. 2023) and SNP-derived genetic relatedness data (Shadrin et al. 2021; van der Meer et al. 2021) in multiple ways. First, whereas prior heritability estimates used region-of-interest level data(Maes et al. 2023), this analysis represents a mass univariate approach in which separate models were run for each vertex of the cortical surface, allowing for the estimation of random effects at a more granular level. This fine-grained analysis led to the observation of heterogeneous effects even within individual cortical regions. Second, while twin heritability studies typically estimate variance components attributable to additive genetic relatedness and shared family, this analysis leverages the ABCD Study^®^ longitudinal data to estimate an additional variance component that is attributable to subject-specific variance, i.e., the variance that is stable within a given participant over multiple study visits. This *S* component, which can be considered a measure of test-retest reliability independent of genetic heritability, accounted for a large proportion of phenotypic variance in all cortical phenotypes studied, representing that this phenotype has a large proportion of variance that is *not* attributable to genetics, common family environment, or the fixed effects, but nevertheless remains stable in a given participant.

Compared to a recent ROI-based study using the ABCD Study^®^ twin sample (Maes et al. 2023), our *A* estimates were smaller, though larger than single nucleotide polymorphism (SNP) heritability estimates derived from GWAS (Shadrin et al. 2021; van der Meer et al. 2021). This is consistent with prior comparisons of twin versus non-twin analyses in the ABCD Study^®^ (Smith et al. 2023). Notably, compared to classical twin studies, the present analysis incorporated the full ABCD Study^®^ sample including siblings and unrelated participants, which may lead to narrower confidence intervals when estimating heritability (Smith et al. 2023). Consistent with previous findings that heritability estimates tend to be lower for phenotypes measured at the regional level compared to global metrics (Maes et al. 2023), and vertex-wise heritability estimates can differ from ROI-based estimates (Eyler et al. 2012), the present study provides evidence that genetic contributions to cortical structure can vary continuously even within a region of the cortex. In addition, the vertex-level distribution of random effects within ROIs confirms that traditional anatomical parcellations do not necessarily match genetic parcellations (Chen et al. 2012, 2013). For example, the supramarginal cortex represented an area with widely distributed effects estimates for all three phenotypes studied, indicating that this region of the cortex may contain several subregions with different amounts of genetic and environmental influence.

This analysis also extends prior heritability estimation by employing a model that incorporated the random effect of subject *S*. Compared to classical twin models that include additive genetic variance, common environment, and one *E* component encompassing both unique environmental influences and measurement error (see Neale and Maes 2004), this analysis allows us to partition unshared variance between siblings into portions that are stable within participants (analogous to test-retest reliability) versus an *E* component that in our model reflects the unshared variance that is *not* stable – potentially representing change within a participant over time and/or measurement error. Importantly, because *S* was included in a model that also contains additive genetic contributions (*A*), the *S* component represents variance that differs between identical twins while remaining stable for a given participant over time, akin to a “cortical fingerprint”. This variance component was large in several distinct regions of the brain for each phenotype assessed, indicating that similar to genetic heritability (Panizzon et al. 2009), subject-specific variance has differential influences on cortical surface area, cortical thickness, and sulcal depth. The *S* component for cortical surface area was particularly strong in a portion of the occipitoparietal cortex that included parts of the supramarginal, superior parietal and inferior parietal cortices. On the other hand, the *S* component for cortical thickness was largest in parts of the cingulate cortex, superior parietal cortex, and the precuneus. Notably, sulcal depth exhibited very large *S* estimates globally with few exceptions. This “fingerprint”-like phenomenon is particularly interesting given the associations that have previously been found between sulcal depth and aging (Rettmann et al. 2006) as well as mental health (Shin et al. 2022). These results imply that all three phenotypes, and particularly sulcal depth, are subject to a substantial amount of influence from environmental factors that are not shared among twins or siblings, which may include random influences during development or unique experiences that influence individual patterns of cortical maturation (Tamnes et al. 2017).

The present study confirms and extends prior literature estimating heritability of cortical phenotypes in adolescents (Maes et al. 2023), using a novel statistical package to leverage the full ABCD Study^®^ sample across multiple timepoints. The results of this large-scale analysis are intended to provide benchmarks for the vertex-wise estimation of variance components including not only the standard ACE model but also the subject-specific variance component, which provides and estimate for the component of test-retest reliability that is not accounted for by shared genetics and family environment. The ABCD Study^®^ cohort is a diverse adolescent sample that was recruited to reflect the adolescent population in the United States; it is possible that the results of this analysis may not generalize to populations with different genetic and cultural backgrounds (e.g., Fan et al. 2023). In addition, while the mixed-effects models used in this work remain agnostic regarding the specific sources of genetic influence, future work should aim to incorporate genomic data to better understand the specific longitudinal contributions of genetic variation to adolescent brain development.

## Acknowledgements

Data used in the preparation of this article were obtained from the Adolescent Brain Cognitive Development^SM^ (ABCD) Study (https://abcdstudy.org), held in the NIMH Data Archive (NDA). This is a multisite, longitudinal study designed to recruit more than 10,000 children age 9-10 and follow them over 10 years into early adulthood. The ABCD Study® is supported by the National Institutes of Health and additional federal partners under award numbers U01DA041048, U01DA050989, U01DA051016, U01DA041022, U01DA051018, U01DA051037, U01DA050987, U01DA041174, U01DA041106, U01DA041117, U01DA041028, U01DA041134, U01DA050988, U01DA051039, U01DA041156, U01DA041025, U01DA041120, U01DA051038, U01DA041148, U01DA041093, U01DA041089, U24DA041123, U24DA041147. A full list of supporters is available at https://abcdstudy.org/federal-partners.html. A listing of participating sites and a complete listing of the study investigators can be found at https://abcdstudy.org/consortium_members/. ABCD consortium investigators designed and implemented the study and/or provided data but did not necessarily participate in the analysis or writing of this report. This manuscript reflects the views of the authors and may not reflect the opinions or views of the NIH or ABCD consortium investigators.

The ABCD data repository grows and changes over time. The ABCD data used in this report came from ABCD Study Data Release 4.0 (http://dx.doi.org/10.15154/1523041). DOIs can be found at https://nda.nih.gov/abcd/abcd-annual-releases.html.

The authors wish to thank the youth and families participating in the ABCD Study^®^ and all staff involved in data collection and curation. The work was also funded by the European Union’s Horizon 2020 research and innovation programme (grant agreement number *964874* to Dr Andreassen, Marie Skłodowska-Curie grant agreement number 801133 to Dr Parekh) and from the Research Council of Norway (*#*223273, # 324252, *#324499)*. D. Smith was supported by Kavli Innovative Research Grant under award number 2022-2195 for this project.

## Author Contributions

Conceptualization, D.M.S., T.E.N., T.L.J., and A.M.D.; Methodology, D.M.S., T.E.N., and A.M.D.; Software, D.M.S., P.P., O.F., and A.M.D.; Formal Analysis, D.M.S.; Resources, T.L.J. and A.M.D.; Writing – Original Draft, D.M.S. and J.K.; Writing – Review & Editing, D.M.S., P.P., R.L., O.F., T.E.N., O.A.A., and T.L.J.; Visualization, D.M.S.; Supervision, O.A.A., T.L.J., and A.M.D.; Funding Acquisition, D.M.S., P.P., O.A.A., T.L.J., and A.M.D.

## Data availability

ABCD Study^®^ data release 4.0 is available for approved researchers in NIMH Data Archive (NDA; http://dx.doi.org/10.15154/1523041).

## Code availability

The code to analyze the data and generate all figures of this manuscript is available on GitHub: https://github.com/dmysmith/random_effects

