## Supplementary Information for "Partitioning variance in cortical morphometry into genetic, environmental, and subject-specific components"

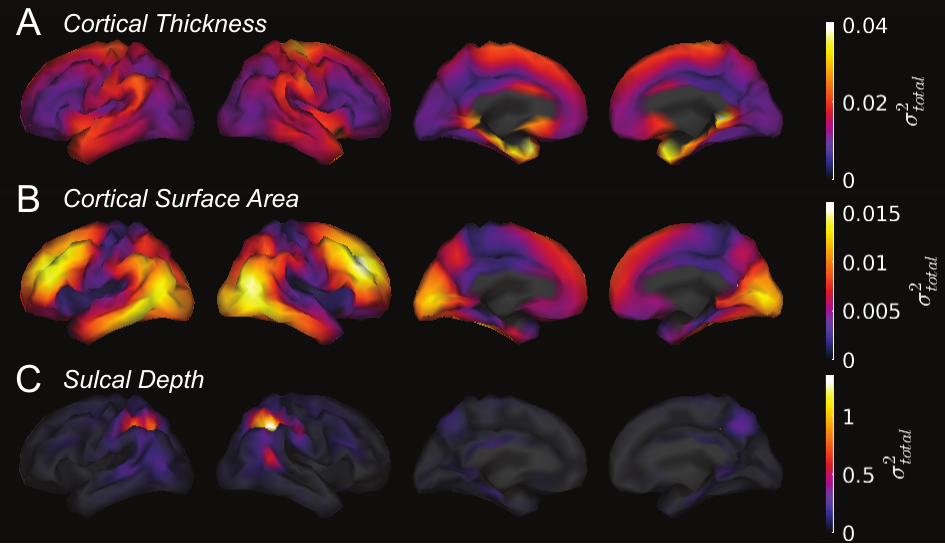

**Supplementary Figure 1.** Vertex-wise maps of total residual variance in cortical thickness (A), cortical surface area (B), and sulcal depth (C). ${\sigma^{2}}_{total}$: Total residual variance after accounting for fixed effects of age, sex, MRI scanner, and software version. ${\sigma^{2}}_{total}$ is expressed in units that match the units of the phenotype of interest (mm for cortical thickness, mm^2^ for cortical surface area, mm for sulcal depth) and represents the total phenotypic variance that is then partitioned into A, C, S, and E components.

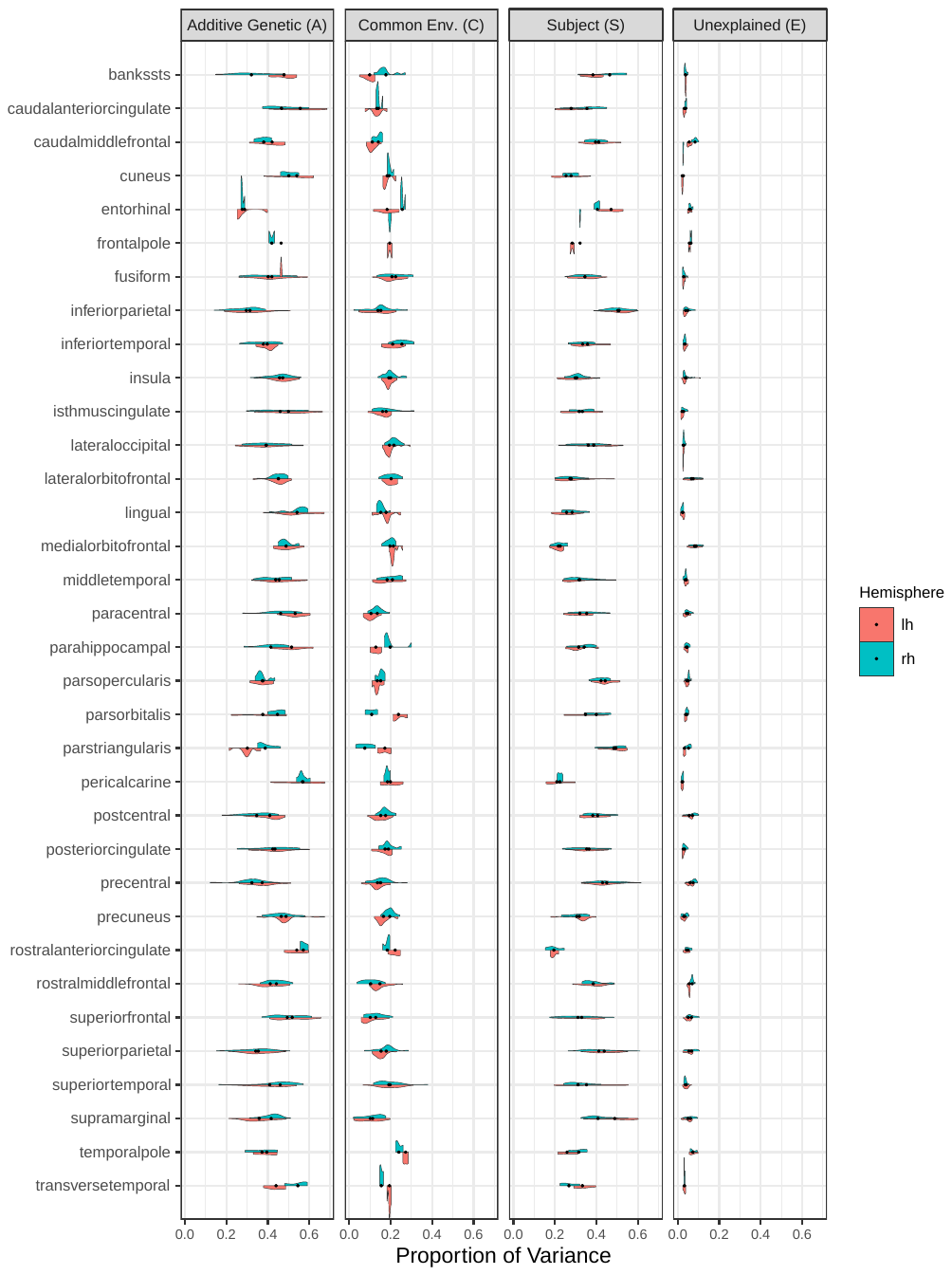

**Supplementary Figure 2.** Vertex-wise additive genetic (A), common environment (C), subject-specific (S), and unexplained (E) variance components in cortical surface area, categorized into regions of interest using the Desikan-Kiliany parcellation and stratified by hemisphere. Marked points represent the mean random effect estimate across all vertices in a given region.

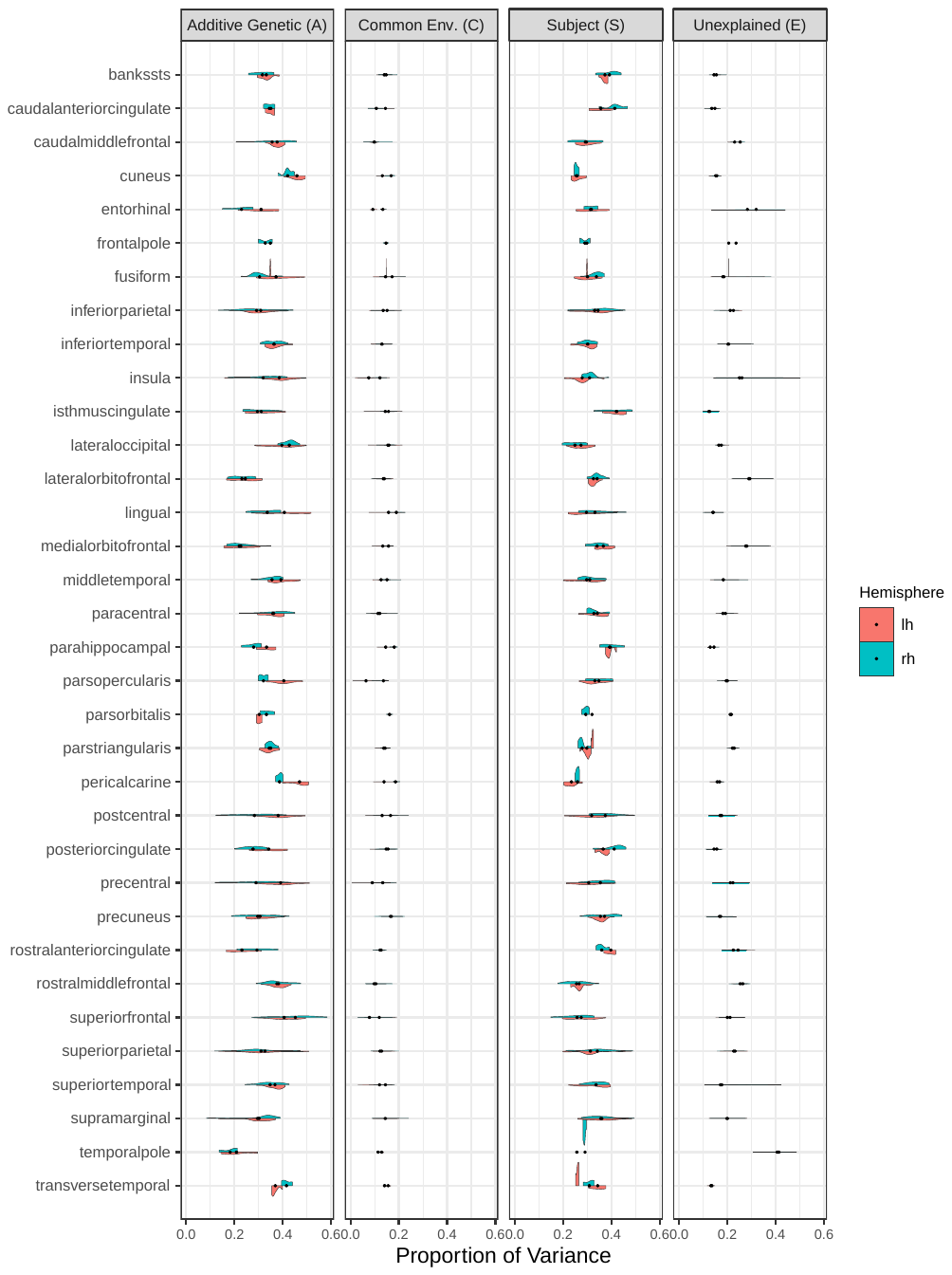

**Supplementary Figure 3.** Vertex-wise additive genetic (A), common environment (C), subject-specific (S), and unexplained (E) variance components in cortical thickness, categorized into regions of interest using the Desikan-Kiliany parcellation and stratified by hemisphere. Marked points represent the mean random effect estimate across all vertices in a given region.

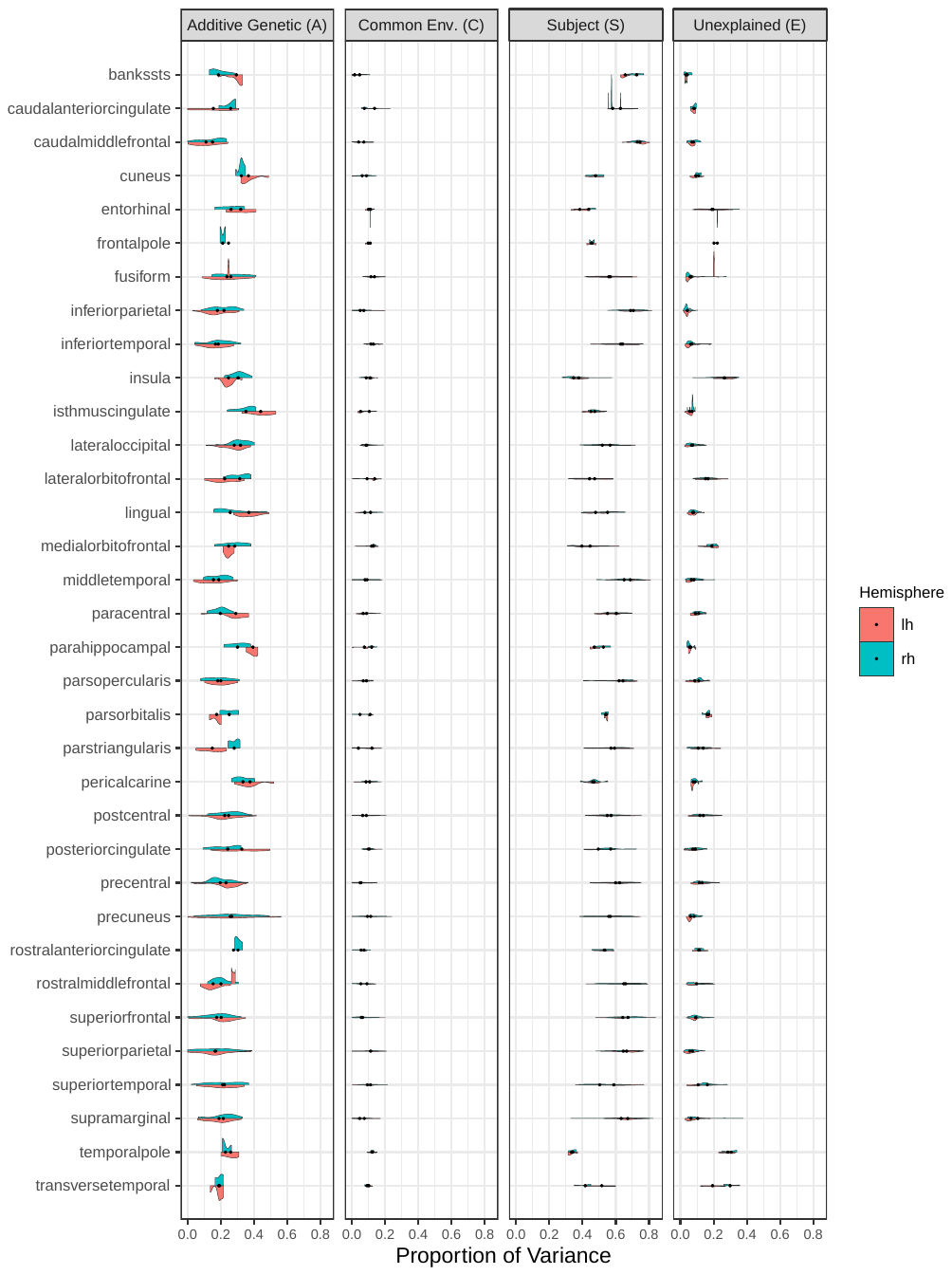

**Supplementary Figure 4.** Vertex-wise additive genetic (A), common environment (C), subject-specific (S), and unexplained (E) variance components in sulcal depth, categorized into regions of interest using the Desikan-Kiliany parcellation and stratified by hemisphere. Marked points represent the mean random effect estimate across all vertices in a given region.

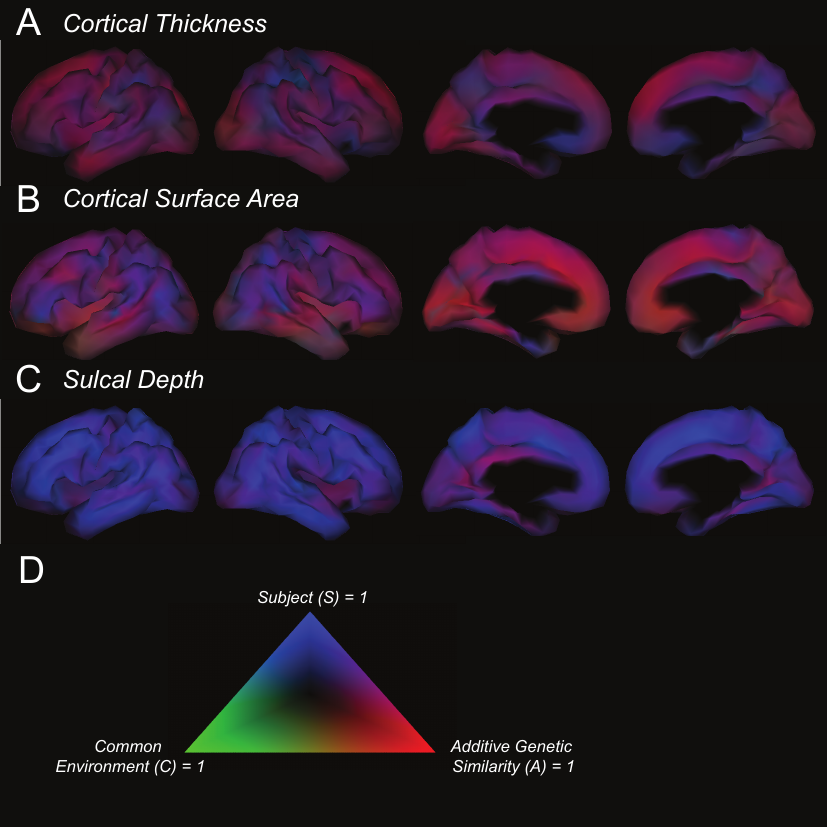

**Supplementary Figure 5.** Additive genetic, common environment, and subject-specific variance component estimates in cortical thickness **(A)**, cortical surface area **(B)**, and sulcal depth **(C)**, visualized on red-green-blue color scale as described in legend **(D)**. Cortical thickness and cortical surface area display widespread patterns in which the majority of variance is explained by additive genetic similarity and subject specific variance, whereas most variance in sulcal depth is explained by subject-specific variance with a lower proportion explained by additive genetic similarity. None of the phenotypes exhibit substantial components of variance attributable to common family environment.

**A**

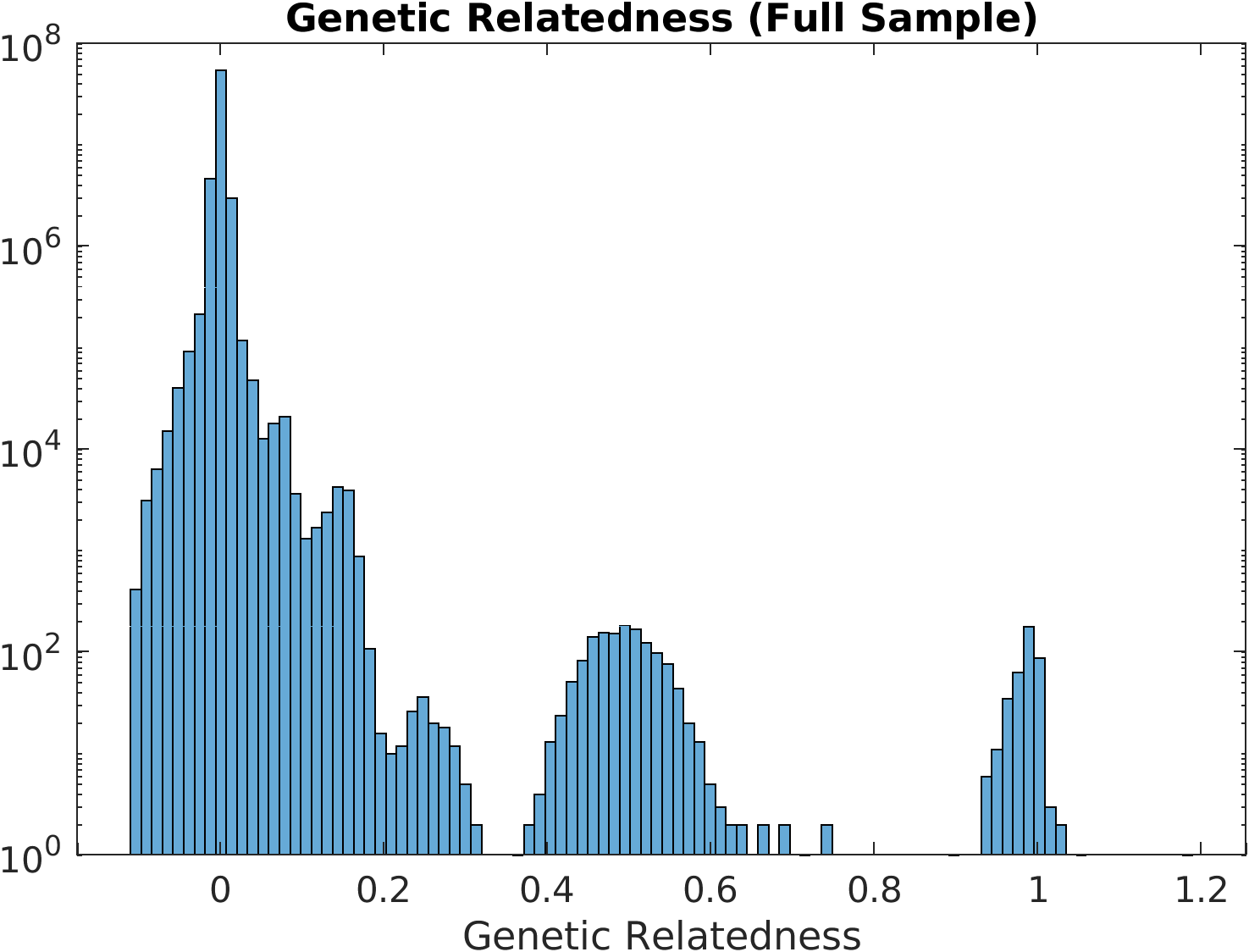

**B**

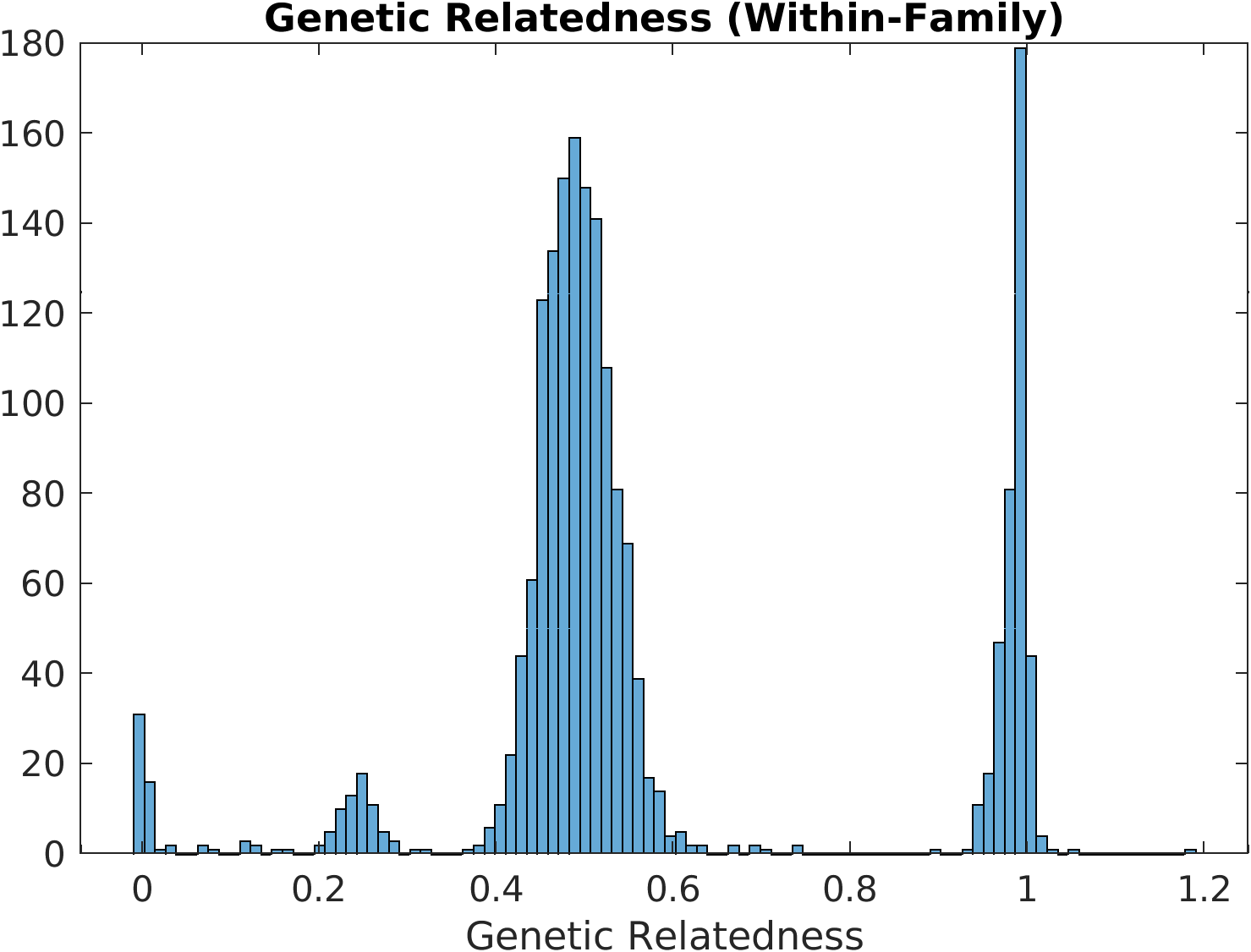

**Supplementary Figure 6.** Genetic relatedness between all pairs in the full analytic sample (A) and the subsample of pairs who share family ID (B).
